## Supplementary figures and images for "From Single Neurons to Behavior in the Jellyfish *Aurelia aurita*"

### Animation of MNN activity and the corresponding swimming motion

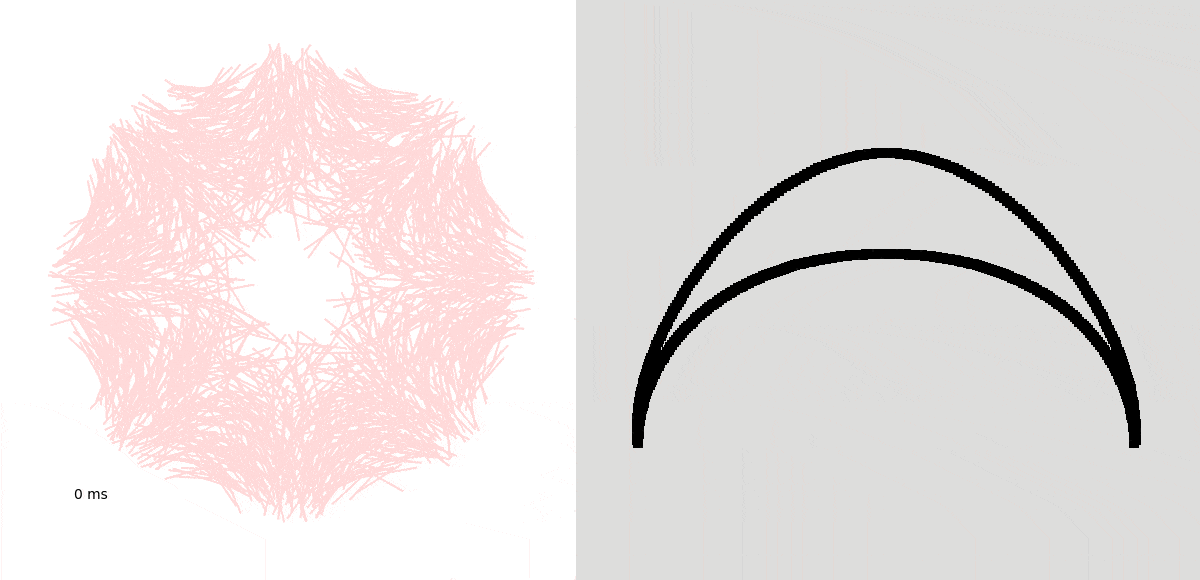

### Supplement Fig. 4

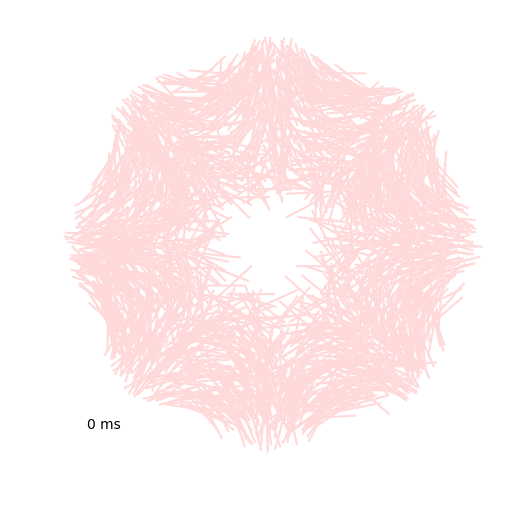

### Supplement Fig. 6

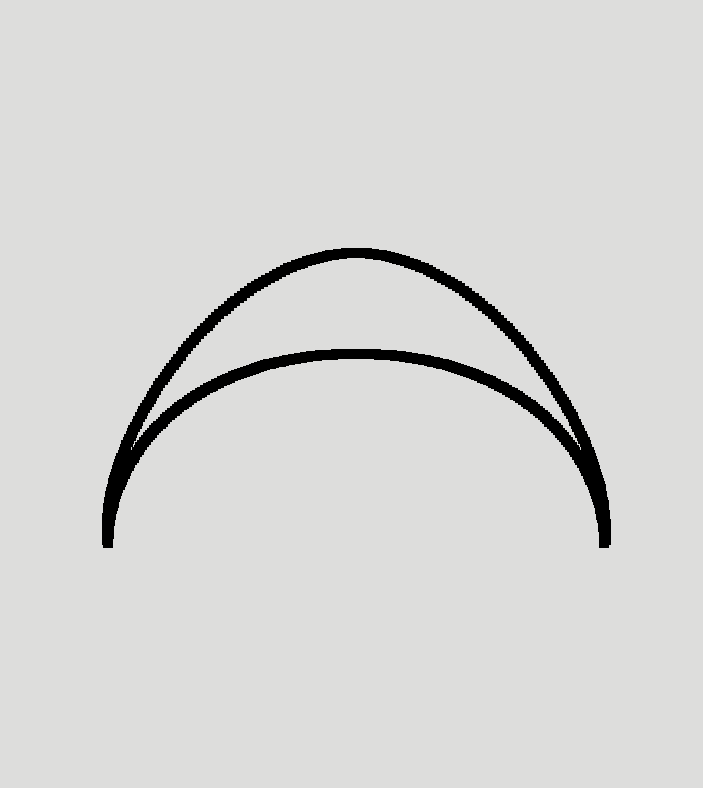

### Supplement Fig. 9

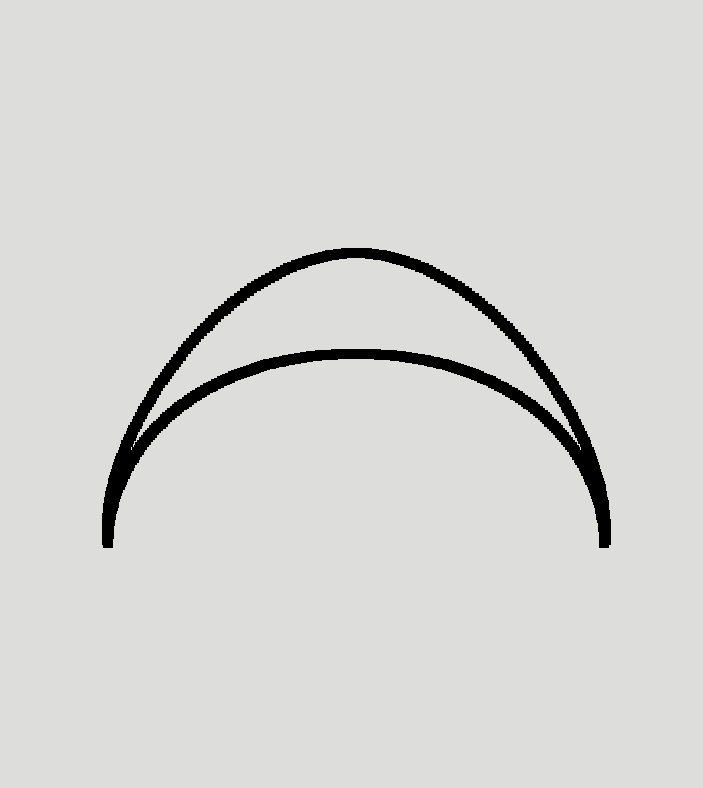

### Supplement Fig. 11

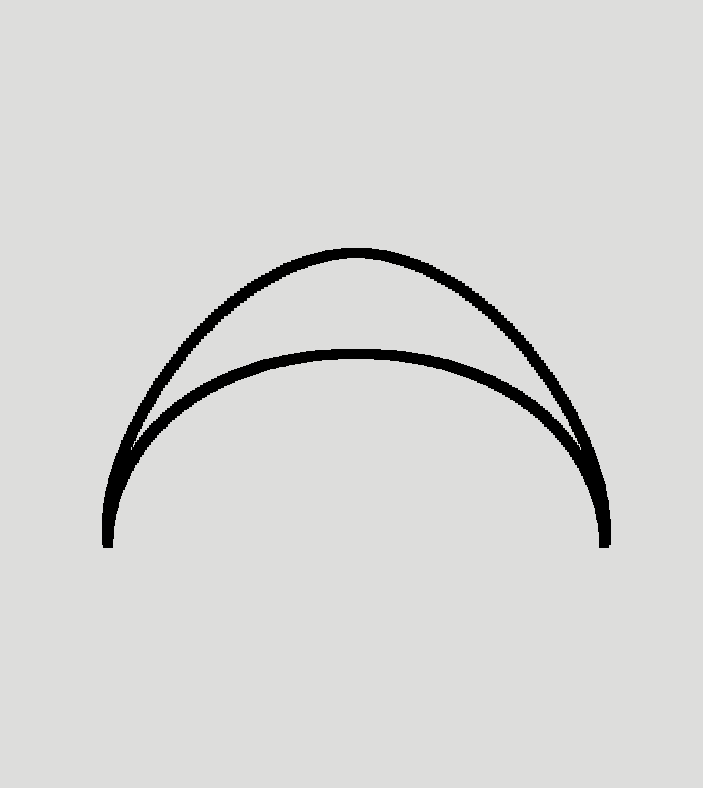
